# PHYLOGENETIC SUPPORT FOR THE EVOLUTION OF APOSEMATISM BEFORE GREGARIOUSNESS IN CONIFER SAWFLIES

**DOI:** 10.64898/2026.11.10.765549

**Authors:** Katja Koskenpato, Philip A. Downing, Saskia Wutke, Catherine R. Linnen, Heikki Helanterä, Marko Mutanen, Carita Lindstedt

**Affiliations:** Faculty of Agriculture and Forestry, Department of Forest Sciences, University of Helsinki, Helsinki, Finland; Ecology and Genetics Research Unit, University of Oulu, Oulu, Finland; Animal Ecology, Department of Ecology and Genetics, Uppsala University, Uppsala, Sweden; Department of Environmental and Biological Sciences, University of Eastern Finland, Joensuu, Finland; Department of Biology, University of Kentucky, Lexington, Kentucky, USA

**Keywords:** conifer sawflies, initial evolution of aposematism, phylogeny, social behaviour

## Abstract

The evolutionary origin of aposematism remains a long-standing puzzle: how can a conspicuous mutant persist and spread within a cryptic prey population? Two competing hypotheses have been proposed. The group-first hypothesis suggests that aposematism evolved secondarily in already aggregating, chemically defended cryptic ancestors, whereas the aposematism-first hypothesis suggests that aposematism arose in solitary ancestors, with gregariousness evolving subsequently. Currently, these two evolutionary pathways have only been studied in caterpillars of Lepidopteran species, but we lack wider taxonimical evidence especially from systems with cryptic but chemically defended species. We evaluated these hypotheses using phylogenetic reconstructions in chemically defended conifer sawflies (Hymenoptera: Diprionidae), a group whose larvae vary in both colouration and aggregation behaviour. Almost half the species examined (20/45) had aposematic and gregarious larvae. Aposematism and gregariousness were positively evolutionarily correlated (*r* = 0.29). Aposematic and gregarious species tended to evolve from aposematic, solitary ancestors, while species with cryptic and gregarious larvae (14/45) tended to evolve from aposematic and gregarious ancestors, a pattern consistent with the aposematism-first hypothesis.

## INTRODUCTION

Aposematic species display conspicuous warning signals, such as orange, yellow, red and white colour patterns combined with the black pattern elements. These signals are used to communicate the presence of unpalatable traits such as chemical defences, spines, or hairs to predators. Predators then learn to associate the warning signal with the unpleasant experience and avoid attacking similar looking individuals in future encounters (avoidance learning). As Wallace put it, the prey must indicate it is a “disgusting morsel” (Darwin 1871). It is well established both theoretically (Leimar et al. 1986, Endler & Mappes 2004) and empirically (Alatalo & Mappes 1996, Lindström et al. 1999, Riipi et al. 2001) that more conspicuous warning signals are easier for predators to learn and remember in comparison to weak visual signals. Avoidance learning becomes even more effective when aposematic prey are aggregated in groups (Alatalo & Mappes 1996, Riipi et al. 2001, Beatty et al. 2005), as predators encounter the aposematic prey more frequently.

However, the origin of aposematism is more challenging to explain (Alatalo & Mappes 1996, Tullberg et al. 2000). This is because a rare, aposematic mutant in a cryptic population should have an increased predation risk due to increased detectability to naïve predators (i.e., predators that have no prior experience in aposematic prey) making it difficult for an aposematic allele to spread and become abundant (Mappes et al. 2005, Ruxton & Sherratt 2006). Hence, aposematic signalling has no value until predators associate it with unpalatability. There are two main hypotheses for how organisms escape this conundrum, allowing aposematism to evolve. These differ in what came first: grouping behaviour (group-first hypothesis) or warning colouration (aposematism-first hypothesis).

The group-first hypothesis proposes that aposematism evolves within kin aggregations of chemically defended, but cryptic prey; even if an aposematic individual is killed, alleles for the warning signal remain in the population via their kin (Fisher 1930, Guilford 1985, Best et al. 2018). However, experimental studies with naïve predators suggest that the group-first hypothesis does not require kin-selection if there are direct benefits of gregariousness (Alatalo & Mappes 1996, Riipi et al, 2001). For example, large group size itself can act as signal or benefit chemically defended prey via dilution effects (Riipi et al. 2001, Speed & Ruxton 2014); if the predator cannot eat the whole group, an individuals’ risk of being eaten is lower in a larger than a smaller group (Riipi et al. 2001, Hunter 2000, but see Lindstedt et al. 2006; 2011). This could be due to predator satiation and/or accumulation of toxins (Skelhorn & Rowe 2007).

The aposematism-first hypothesis is based on signal augmentation theory (Gamberale & Tullberg 1996, Riipi et al. 2001) and suggests the opposite pathway to the group-first hypothesis. Under this hypothesis, conspicuous warning colouration is proposed to evolve first in solitary individuals and works because predators are neophobic or conservative, preventing them from attacking novel conspicuously coloured prey (Mallet & Singer 1987, Marples & Kelly 1999, Marples et al. 2005, Mappes et al. 2005). Alternatively, aposematic prey with good secondary defences can survive an attack without lethal injuries (Wiklund & Järvi 1982). Once warning colouration is established, gregariousness is expected to follow because it strengthens the signal (Gamberale & Tullberg 1996) and speeds up avoidance learning (Alatalo & Mappes 1996, Riipi et al. 2001, Beatty et al. 2005, Sillén-Tullberg 1990 but see Tullberg et al. 2000). Gregariousness could also be favored via other benefits such as more effective exploitation of the host plant (Ghent 1960, Fordyce & Nice 2004, Reader & Hochuli 2023) or maintenance of optimal microclimatic conditions (Seymour 1974, Despland 2021, Ostwald et al. 2022, Qian et al. 2024). These other potential benefits of gregariousness are not mutually exclusive with improved warning signalling.

Comparative studies of Lepidopteran caterpillars support the aposematism-first hypothesis, finding that the evolution of secondary defences and aposematism often preceded the evolution of gregariousness (Sillén-Tullberg 1988, Tullberg & Hunter 1996, Beltran et al. 2007, McLellan et al. 2023). Recent comparative work further indicates that the ancestral Lepidopteran caterpillar was likely aposematic and solitary, and that colouration might change more easily than gregariousness (Wang et al. 2021). However, these studies could not measure chemical defence directly and assumed that warning colouration means unpalatable and cryptic means palatable (Sillén-Tullberg 1988, McLellan et al. 2023). Under this assumption, a cryptic, chemically defended and gregarious species is scored as palatable. Therefore, these data sets do not allow us to test whether gregariousness in chemically defended prey preceded aposematism, which has been the basic assumption behind the “group-first” hypotheses (Ruxton & Sherratt 2006).

Furthermore, currently most of the evidence comes from Lepidopterans, but we do not know if the observed patterns are generalizable to other taxonomic groups. For example, in the aposematism-first model, the likelihood of aposematic prey surviving or escaping an attack by a predator may also depend on prey type and how chemical defences are displayed (Wiklund & Järvi 1982). Therefore, evaluating the relative contributions of “aposematism-first” vs. “group-first” scenarios to the diversity of anti-predator strategies observed in nature requires reconstructing the order in which conspicuousness and grouping evolved across a wider range of taxa.

Conifer sawflies (Hymenoptera: Diprionidae) allow us to address both gaps. We investigated the evolution of aposematism and gregariousness in diprionid larvae. Diprionidae comprise roughly 140 described species across 11 extant genera (Wagner & Raffa 1993, Taeger et al. 2018, Davis et al. 2023). These species vary extensively in terms of their protective colouration (ranging from cryptic to aposematic) as well as levels of gregariousness, ranging from solitary to large groups of 30-100 or more individuals (Figure 1, Terbot et al. 2017, Davis et al. 2023, Lindstedt et al. 2025, Ritter et al. 2026). All diprionid larvae are assumed chemically defended (Davies et al. 2023). Thus, there are species that are chemically defended yet cryptic. Larvae feed on conifers in the families Pinaceae (subfamily Diprioninae) and Cupressaceae (subfamily Monoctinae) and sequester host-plant resins for their chemical defence (Eisner et al. 1974, Björkman & Larsson 1991, Codella & Raffa 1995, Davies et al. 2023).

**Figure 1.**
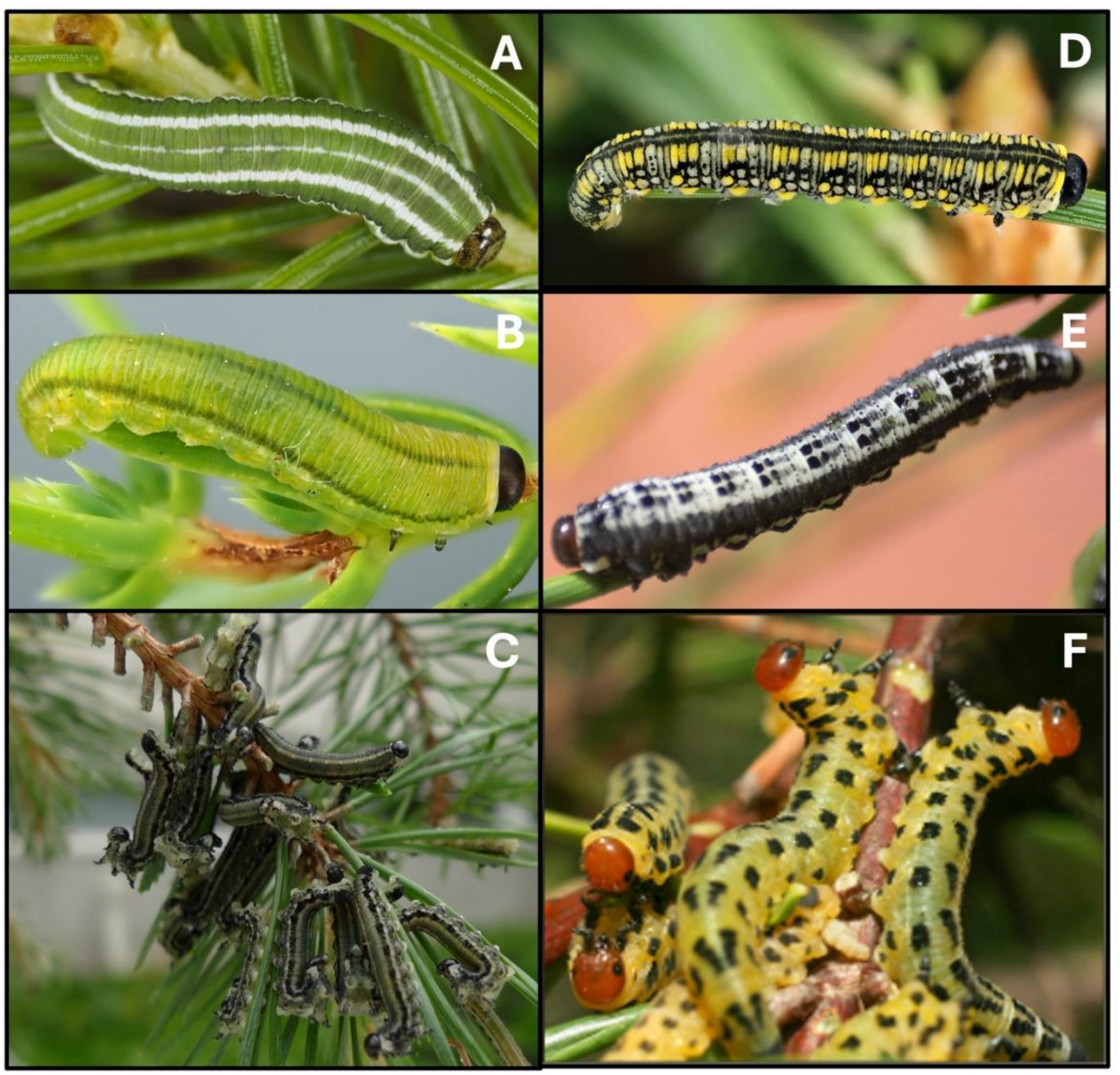
Variation in gregariousness and colouration in diprionid larvae: cryptic and solitary *Gilpinia hercyniae* (A), cryptic and solitary *Monoctenus juniperi* (B), cryptic and gregarious *Neodiprion sertifer* taking the U-posture and producing defensive fluid (C), aposematic and solitary *Diprion similis* (D), aposematic and gregarious *Gilpinia socia* (E) and aposematic and gregarious *Neodiprion lecontei* (F). Photographs: Insect Bioliteracy Group, University of Oulu (A & B), Carita Lindstedt (C), Mikko Vallinmäki (D), Benny Hjelt (E), Robin Bagley (F).

When diprionid larvae are attacked, they take a defensive U-bend shaped posture and deploy resinous defensive fluid from their mouths (Eisner et al. 1974). If the predator comes closer, the larvae try to actively direct the fluid towards the predator or parasitoid (Eisner et al. 1974). The chemical defence of diprionids deters a range of predators such as birds, spiders and ants (Eisner et al. 1974, Sillén-Tullberg 1990, Codella & Raffa 1996, Lindstedt et al. 2006; 2011). Diprionid larvae also cooperate in chemical defence, gaining both direct and indirect (improved survival of their kin) fitness benefits from contributing to collective defence (Ritter et al. 2026, Van Meyel et al. 2026).

In general, the responses of predators support key assumptions of warning signal theory. While chemical defence in itself is enough to educate predators to avoid cryptically coloured pine sawfly larvae (Sillén-Tullberg 1990, Ritter et al. 2026, Van Meyel et al. 2026), avoidance by predators becomes more effective when pine sawfly larvae are in groups (Sillén-Tullberg 1990, Van Meyel et al. 2026) or if they display traditional aposematic yellow-black or white-black warning colouration (Linnen et al. 2018, Lindstedt et al. 2022). Thus, both gregariousness and warning colouration improve avoidance of chemically defended larvae, so both orders of evolution are plausible. Determining which trait arose first in diprionids requires a phylogenetic reconstruction.

Diprionidae offer an ideal system for studying the evolutionary interplay between aposematism and gregariousness because other traits vary liitle among species. All Diprionidae species share a similar visual background (spending the larval stage feeding on conifer needles), a broadly similar chemical defence profile and responsive defence behaviour (Davis et al. 2023). To this end, we ask: (1) do gregariousness and aposematism evolve in a correlated fashion, and if so, (2) is there a tendency for changes in one trait to occur before changes in the other? To answer these questions, we present a new comprehensive Diprionid phylogeny (N = 58 species, covering 9 of 11 genera and the full range of colouration and group sizes) based on molecular data and use it to analyse how colouration and gregariousness have evolved in Diprionidae.

## MATERIALS AND METHODS

To test for the correlated evolution of aposematism and gregariousness in Diprionidae, we conducted a phylogenetic comparative analysis. This involved four main steps: (1) collecting data from the published literature on gregariousness for as many diprionid species as possible, (2) categorizing larval colouration (cryptic or aposematic), (3) building a phylogenetic tree using molecular methods, and (4) using two statistical approaches to investigate evolutionary relationship between traits.

### Trait data collection

Data on gregariousness was collected from research papers, review papers, books, leaflets and webpages. Searches were done in Google by typing in all scientific species names separately. Data were collected between 11/2022 – 9/2026. Gregariousness was scored as a binary variable, where gregariousness was either present (species is gregarious) or absent (species is solitary). We gathered trait data only from mid- to late-feedining instars as the colouration and behaviour can change drastically with developmental stage (Herrig et al. 2021, Davis et al. 2023).

### Colour scoring

We obtained pictures of diprionid larvae from iNaturalist, webpages, articles and from the collections of MM and CRL. Species identifications and developmental stages were confirmed by CRL and MM. Larvae were categorised as either aposematic or cryptic (binary variable). If larvae had clear components (e.g., dots or stripes) of black on a otherwise light (white or yellow) body making clear contrasts, the species were categorised as aposematic. Larvae not showing such colour patterns (in most cases uniform greenish-brownish colouration and some cases with longitudal stripes) were categorized as cryptic. Scoring of all the species was done by CL, KK, CRL and MM and followed the same logic and binary classification as used in previous studies with Lepidopteran caterpillars (Sillén-Tullberg 1988, Tullberg & Hunter 1996, McLellan et al. 2023).

### Phylogenetic trees

The library preparation, high-throughput sequencing and assembly of ultraconserved elements (UCEs) followed the procedures described in Wutke et al. (2024). In short, we followed the PHYLUCE pipeline (Faircloth 2016) with default settings for generating UCE assemblies and alignments. For the phylogenetic analyses, we used a 75 % complete data matrix (i.e., based on loci present in at least 75 % of the taxa, alignment length 397,119 bp, concatenated from 1279 UCE loci). The maximum likelihood (ML) phylogeny was inferred using the MPI version of RAxML 8.2.12 (Stamatakis 2014), applying the rapid bootstrap algorithm (-f a), 1000 bootstrap replicates, and GTRGAMMA site rate substitution model. The resulting phylogenetic tree was rooted using *Augomonoctenus smithii* as an outgroup, as previous analyses suggest this taxon as sister to the remaining Diprionidae (Wutke et al. 2024). Divergence dating was conducted in BEAST v2.7.7 (Bouckaert et al. 2014) using the rooted ML tree as fixed topology and *Eodiprion groehni* as fossil calibration. We further used a uniform root prior, an uncorrelated relaxed lognormal clock model, and birth–death tree prior. Run convergence was confirmed in Tracer v1.7.1 (Rambaut et al. 2018) by checking that effective sample sizes (ESS) were *>* 200. The maximum clade credibility (MCC) tree was summarized from the combined runs with LogCombiner v2.6.3 and TreeAnnotator v2.6.3 after discarding the initial 10 % of trees as burn-in. We also ran the analysis without data by sampling only from the prior.

### Statistical analyses

Two methods were used to investigate the evolutionary association between gregariousness and aposematism. First, we used the threshold model (Felsenstein 2012, Revell 2014) to estimate the evolutionary correlation between aposematism and gregariousness. This model works by assuming that each binary trait is underlain by an unobserved, normally distributed continuous trait, called the liability. The binary trait changes state when the liability crosses a certain threshold value. These thresholds are unknown and are simulated onto a phylogeny following Brownian motion. We then translate the liabilities back into binary traits and compute the likelihood of the model based on the match between the simulated and observed distribution of binary character states at the tips of the phylogeny. The evolutionary correlation between binary traits is simply the correlation of their underlying liabilities. In our case, a positive correlation would indicate that lineages evolving aposematism tend to evolve gregariousness, and vice versa. Importantly, this approach measures association rather than causation: it cannot tell us if one trait is a necessary precursor for the other. The threshold model was fit using the *phytools* R package (Revell 2024) and was run for one million generations.

Second, we used a modified form of Pagel’s (1994) method to investigate the order in which aposematism and gregariousness most likely evolved. We scored our two binary traits as a single multi-state character with four possible outcomes: 1 = cryptic and solitary, 2 = cryptic and gregarious, 3 = aposematic and solitary, 4 = aposematic and gregarious. We then constructed three main transition rates matrices corresponding to a null hypothesis (*H0*), the group-first hypothesis (*H1*) and the aposematism-first hypothesis (*H2*), for the evolution of aposematism (Figure 2).

**Figure 2.**
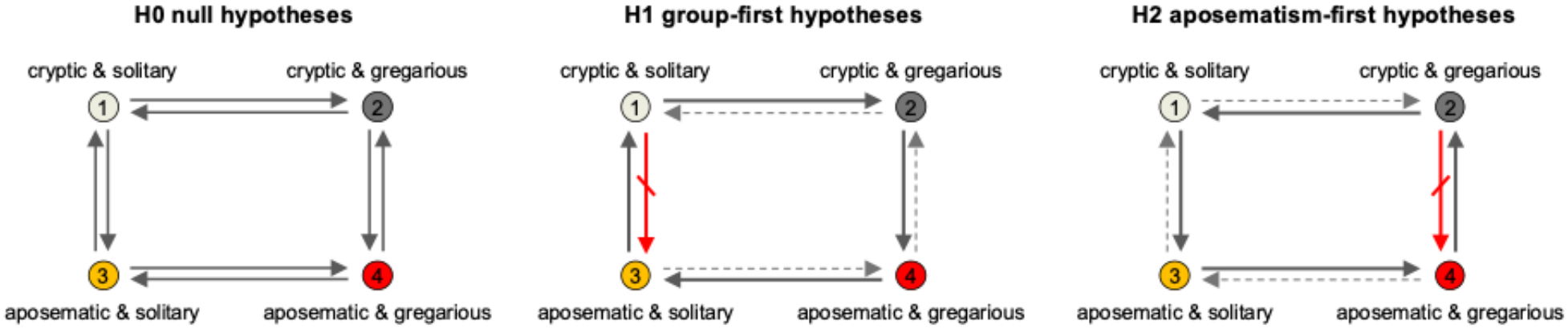
Hypotheses for the evolution of aposematism. In the null hypothesis (*H0*), all changes between states were possible, occurring either at the same (*H0a*) or different rates (*H0b*). In the group-first hypothesis (*H1*), aposematism had to evolve from a gregarious ancestor (solid line, state 2 to state 4) and could not evolve from a solitary ancestor (red line, state 1 to state 3). We allowed unidirectional changes (solid lines) at the same (*H1a*) or different rates (*H1b*) and bidirectional changes at different rates (dashed lines; *H1c*). In the aposematism-first hypothesis (*H2*), aposematism had to evolve from a solitary ancestor (solid line, state 1 to state 3) and could not evolve from a gregarious ancestor (red line, state 2 to state 4). We allowed unidirectional changes (solid lines) at the same (*H2a*) or different rates (*H2b*) and bidirectional changes at different rates (dashed lines; *H2c*).

Our null hypothesis allows aposematism to evolve from a solitary or a gregarious ancestor. We allowed two options: transitions between all states could occur at the same rate (*H0a*), or at different rates (*H0b*). In the group-first hypothesis (H1), aposematism must evolve from a gregarious ancestor (from state 2 to state 4). This means that no transitions from a solitary ancestor to an aposematic descendant are permitted (from state 1 to state 3). We allowed three options: unidirectional changes with transitions between states occurring at the same (*H1a*) or different rates (*H1b*), or bi-directional changes at different rates between states except from state 1 to state 3 (*H1c*). In the aposematism-first hypothesis (H2), aposematism must evolve from a solitary ancestor (from state 1 to state 3) and then gregariousness can evolve (from state 3 to state 4). This means that no transitions from a gregarious ancestor to an aposematic descendant are permitted (from state 2 to state 4). We allowed three options: unidirectional changes with transitions between states occurring at the same (*H2a*) or different rates (*H2b*), or bi-directional changes at different rates between states except from state 2 to state 4 (*H2c*).

These eight models were fit to the data using maximum likelihood in the *corHMM* R package (Boyko & Beaulieu 2021) and compared using AIC. The best fitting model was used to estimate marginal ancestral states, which give the probability that each node is in each state.

## RESULTS

### Phylogenetic relationships and divergence-time estimates

Our phylogenetic reconstruction yielded a generally well-resolved Diprionidae tree, with most nodes receiving strong support (bootstrap support > 95%; Figure 3). The subfamily Diprioninae was recovered as monophyletic. Monocteninae, on the other hand, was rendered paraphyletic by design, as *Augomonoctenus smithii* was selected as the outgroup for rooting the tree. At the genus level, most diprionid genera were recovered as monophyletic, with the exception of *Neodiprion*. This paraphyly arises because *N. xiangyunicus* was placed in a clade alongside *Zadiprion* taxa, rather than clustering with other *Neodiprion* taxa. Within *Neodiprion*, most relationships are consistent with previous phylogenetic work (Linnen & Farrell 2007; 2008a; 2008b, Herrig et al 2024). However, the placement of three species – *N. merkeli*, *N. compar*, and *N. dubiosus* – differed from earlier studies.

**Figure 3.**
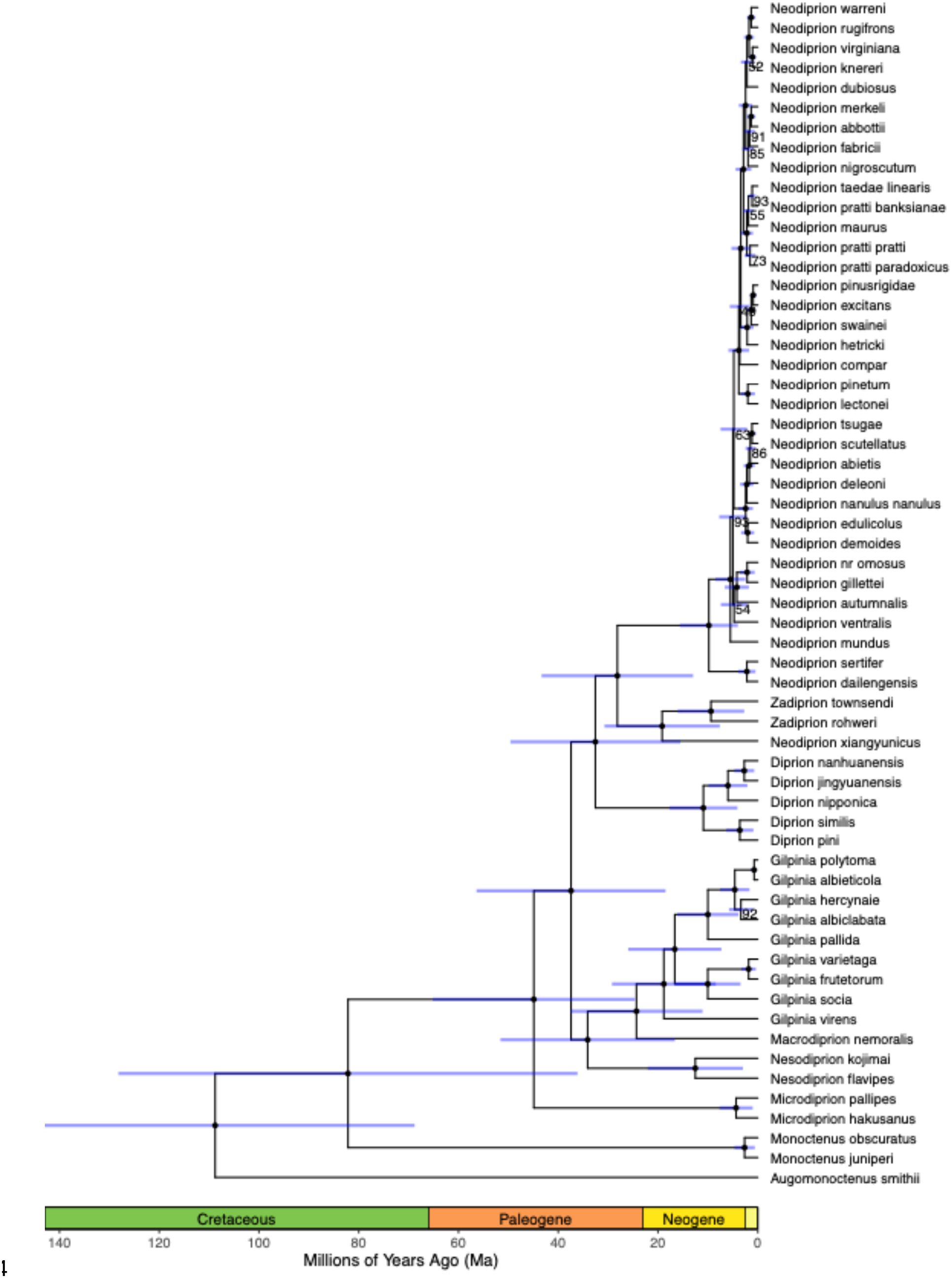
Time-calibrated phylogeny of Diprionidae based on the maximum likelihood topology of the 75 % complete matrix. Blue bars indicate 95% HPD intervals of age estimates. Node support is shown as black dots if bootstrap support (BS) was 95% or above or the actual BS value if below 95%.

### Correlation between gregariousness and aposematism

We successfully collected data on larval colouration and gregariousness for 45 species (out of 58 species in the phylogeny, Table S1). We did not consider subspecies. Three quarters of these species were gregarious (34/45), with 14 of these being cryptic and 20 being aposematic. The remaining 11 species were solitary, with eight of these being cryptic and three being aposematic. Aposematic and gregariousness were positively evolutionarily correlated, indicating that lineages that evolve aposematism tend to evolve gregariousness, although the 95 % confidence interval included zero (*r* = 0.29, 95% CI = -0.25 to 0.76).

### Order of changes in gregariousness and aposematism

Aposematism tended to evolve from solitary rather than gregarious ancestors. These solitary and aposematic ancestors then gave rise to gregariousness and aposematic descendants. Aposematism was then lost from this ancestral state, giving rise to cryptic and gregarious species. These inferences are based on our finding that the aposematism-first hypothesis is best supported by the data: model *H2a* had the lowest AIC scores and accounted for 83% of the weight (Table 1). The group-first hypotheses (*H1*) had the poorest fit to the data, which is particularly evident within the genus *Neodiprion* (Figure 4). Almost all *Neodiprion* species are gregarious (29/30 species), with aposematism as the most likely ancestral state for this clade. In contrast, outside of the *Neodiprion* clade, seven of the eight cryptic species are solitary.

**Figure 4.**
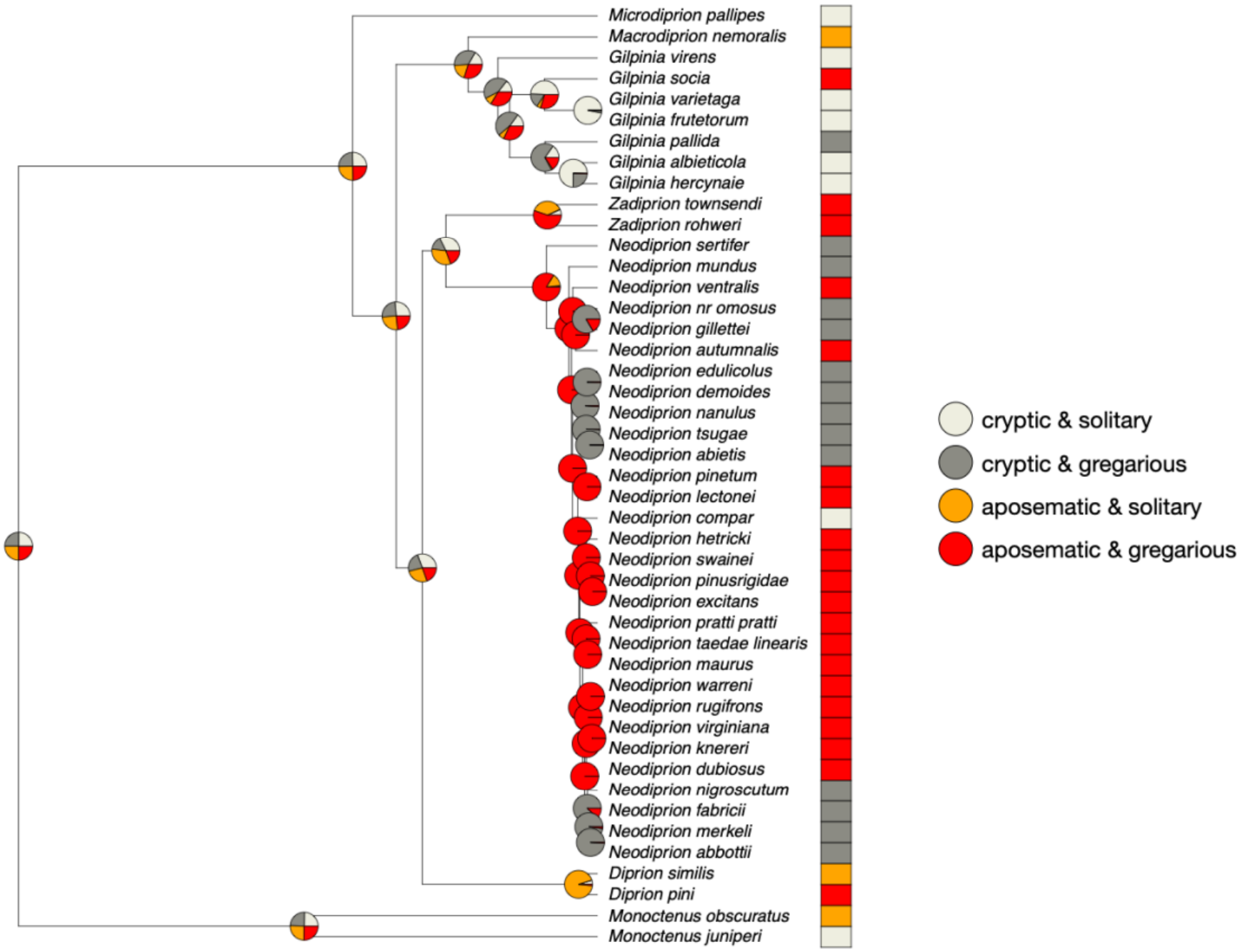
The evolution of larval aposematism and gregariousness across 45 conifer sawfly species. Marginal ancestral states were estimated using the best supported model (*H2a*).

**Table 1.** Log-likelihood and Akaike information criterion (AIC) estimates for the eight models for the evolution of aposematism and gregariousness. Models with the lowest AIC are the best supported and models with an AIC difference < 2 are considered equally good descriptions of the data, while those with a difference of > 5 provide evidence that one is a better fit (Burnham & Anderson 2002).

| **Model** | **log likelihood** | **AIC** | **ΔAIC** | **Akaike weight** |
|---|---|---|---|---|
| *H2a* | -38.7 | 79.4 | 0.0 | 0.83 |
| *H2b* | -38.0 | 84.1 | -4.7 | 0.08 |
| *H0a* | -41.1 | 84.3 | -4.9 | 0.07 |
| *H2c* | -38.0 | 90.1 | -10.7 | 0.00 |
| *H0b* | -38.0 | 92.1 | -12.7 | 0.00 |
| *H1c* | -39.2 | 92.4 | -13.0 | 0.00 |
| *H1b* | -42.2 | 92.4 | -13.0 | 0.00 |
| *H1a* | -44.8 | 91.5 | -12.1 | 0.00 |

## DISCUSSION

We present the most comprehensive molecular phylogenetic tree (58 species) of conifer sawflies (Hymenoptera: Diprionidae) constructed to date. Our results suggest that aposematic colouration is evolutionarily positively correlated with gregariousness: species with aposematic and gregarious larvae made up almost half of our sample (44 %, Figure 4). Ancestral state reconstruction indicates that aposematism most likely evolved from a solitary ancestor and that gregariousness arose subsequently, supporting the aposematism-first hypothesis (Table 1).

In the “group-first” hypothesis, the problem of the initial evolution of aposematism is often presented in a simplified situation where aposematic mutants are assumed to evolve under conditions where predators need to sample, taste, and kill "the first aposematic prey," which is located in an otherwise cryptic prey population. However, Diprionidae are an example of a taxonomic group where individuals display chemical defences externally (as a regurgitated droplet of resin), giving predators a chance to evaluate their unprofitability before actually harming the individual. These responsive chemical defences may have facilitated the early evolution of aposematism in these lineages. Furthermore, it is likely that when the first warning colours evolved in Diprionidae, other aposematic species were already present and local predators were not necessarily unfamiliar with aposematic species and certain signal patterns at the time. This is also supported by experiments with naïve predators: once local predators have encountered aposematic prey, gregariousness is not necessarily needed for the aposematic signals to become more frequent (Alatalo & Mappes 1996, Riipi et al. 2001, Lindström et al. 1999). Thus, gregariousness has not been a necessity for aposematism to become more frequent within the Diprionidae lineage, but it might still play an important role in evolutionarily older lineages.

One reason why the aposematism-first hypothesis received the highest support might be due to the transitions the models for each hypothesis allow. Unlike most group-first models (except for H1c), the aposematism-first models allow direct transition from a solitary, aposematic ancestor to an aposematic, gregarious descendant (state 3 to state 4) i.e., colouration evolves first and gregariousness follows. This pattern is consistent with signal augmentation theory, which posits that warning signals become more effective when encountered more often (Gamberale & Tullberg 1996). In lepidopteran larvae, phylogenetic analyses revealed that transitions from solitary to gregarious occur at higher rate in aposematic lineages than in cryptic ones, and transitions from crypsis to aposematism occur faster in gregarious lineages than in solitary ones (McLellan et al. 2023). The same study found that artificial cryptic larvae suffered from increased predation when grouped, whereas artificial aposematic ones benefitted from groups (McLellan et al. 2023). Supported by previous studies (Gagliardo & Guilford 1993, Alatalo & Mappes 1996, Gamberale & Tullberg 1998, McLellan et al. 2023, Riipi et al. 2001), this suggests that gregariousness and aposematism are mutually reinforcing traits, a pattern we extend to Diprionidae.

While aposematism and gregariousness are evolutionarily correlated, our analyses also suggest that within the *Neodiprion* clade, aposematism was subsequently lost from an already-gregarious ancestor, giving rise to species that are chemically defended, cryptic, and gregarious. This is consistent with phylogenetic patterns observed in amphibians, where warning colouration tends to be evolutionarily transient and is frequently lost (Arbuckle & Speed 2015). There can be several reasons for the loss of aposematism in gregarious lineages in our system. If generalist predators initially avoid aposematic prey but evolve into specialists that can handle those, prey may subsequently re-evolve crypsis (Kikuchi et al.

2021). Additionally, as generalist predators are assumed to maintain aposematism among prey species by avoiding warning-coloured prey, the specialist:generalist predator ratio in the environment may affect the viability of aposematism (Kikuchi et al. 2021). For example, Koskenpato et al. (2026) demonstrated experimentally that increased conspicuousness in the colouration of pine sawfly larvae (*Neodiprion sertifer*) significantly decreases their survival against the natural predator community despite the chemical defences. Therefore, crypsis, combined with gregariousness, can indeed be advantageous in varying environments where predators are naïve or tolerant to prey defences (Alatalo and Mappes 1996, Riipi et al. 2001, Endler & Mappes 2004, Koskenpato et al. 2026). Finally, even though conspicuousness increases the efficacy of warning signals, predators can also learn to avoid chemically defended prey as long as they can distinguish the defended prey from the non-defended prey (Sherratt & Beatty 2003, Merilaita & Ruxton 2007). This can also relax selection for conspicuousness.

Nevertheless, both traits measured here may also provide benefits on their own. Gregariousness is not only an anti-predator strategy, but might be essential in group facilitated feeding (Fordyce & Nice 2004, Reader & Hochuli 2023). For example, small larvae of *Neodiprion pratti* starve when reared alone as they are not able to access the nutritional tissues under the hard needle surface without the help of conspecifics (Ghent 1960). Gregariousness can also improve microclimatic conditions (Despland 2021, Qian et al. 2024). Many Diprionidae females cluster their eggs, which can help in group formation after hatching (Terbot & Linnen 2019, Davis et al. 2023, Lindstedt et al. 2025). However, gregariousness can also be costly as it can lead to e.g. increased resource competition (Despland & Le Huu 2007, Qian et al. 2024) and exposure to predators and pathogens (Anderson & May 1981, Hochberg 1991). As all diprionid larvae are assumed chemically defended, chemical defence appears to be associated with different combinations of defensive colouration and gregariousness in this family. This is consistent with the studies conducted with chemically defended sawfly larvae in the families *Argidae* and *Pergidae*. These lineages show only weak phylogenetic support for any combination of colouration and gregariousness, suggesting that chemical defence alone can result in a range of different trait combinations depending on local predator communities (Boevé et al. 2018).

The reconstruction of the molecular phylogenetic tree resulted in some changes for the placement of species. The genus *Neodiprion* was recovered as paraphyletic due to the placement of *N. xiangyunicus* in a clade together with *Zadiprion* taxa. Since *N. xiangyunicus* and *Zadiprion* occur in non-overlapping geographic regions, sample misidentification is an unlikely explanation for this placement. Instead, this result suggests the need for a focused taxonomic revision of the *N. xiangyunicus* and *Zadiprion* clades: either by reclassifying *N. xiangyunicus* within *Zadiprion* or rather, given the broader pattern, subsuming *Zadiprion* into the *Neodiprion* genus. Within the *Neodiprion* clade, the position of *N. merkeli*, *N. compar* and *N. dubiosus* differed from previous studies. The latter two were particularly unstable: Herrig et al. (2024), which was based on analysis of whole genomes using multiple species-tree methods, similarly found that the placement of *N. compar* and *N. dubiosus* varied depending on the underlying dataset and analytical method used, rather than converging on a single well-supported position. Thus, the differences we observe in the placement of these three species is not surprising: the *Neodiprion* radiation has previously proven difficult to resolve, likely reflecting genuine biological sources of gene-tree heterogeneity (e.g., incomplete lineage sorting and introgression) rather than simply insufficient data (Linnen & Farrell 2007, Herrig et al. 2024). Importantly, regardless of the exact position of these three species, our downstream ancestral state reconstructions are unlikely to be considerably affected.

Several methodological considerations are worth noting when interpreting these results. First, colouration may serve multiple functions depending e.g., on the distance it is viewed (Tullberg et al. 2005; Bohlin et al. 2008; Loeffler-Henry et al. 2023). For example, prey may look cryptic from a distance but aposematic when viewed close by. Colouration may also work in disruptive manner (Stevens et al. 2006, Schaefer & Stobbe 2006). We cannot exclude these potential multiple functions of colouration. Second, colouration and gregariousness can change across larval instars, so we scored colouration and gregariousness consistently from feeding mid-instar larvae. Third, documentation of diprionid larvae is limited, and we could only access data on less than 1/3 of the species. Fourth, we scored colouration by human eye alone and thus may have missed perception by predators that can also e.g., sense ultraviolet light (Cuthill 2006, Kemp et al. 2015).

However, using human perceivers are commonly accepted in similar studies (Sillén-Tullberg 1988, Tullberg & Hunter 1996, Mappes et al. 2014, McLellan et al. 2023) and it is known that ultraviolet signals are rare in pine sawfly larvae (Lindstedt et al. 2022). Fifth, traits were scored as binary variables, although they are inherently more continuous in nature. In addition, chemical defence was not measured at the species level but was assumed to be similar across species based on existing knowledge (Davies et al. 2023). Finally, some diprionid species, e.g. *Neodiprion lecontei*, show intraspecific colour polymorphism as larvae (Linnen et al. 2018, Lindstedt et al. 2022), which is not taken into account in this study. For a better understanding of the correlation between colouration and gregariousness, more data on intraspecific polymorphism is required. However, we are confident that these constraints do not bias our results or their interpretation, as we managed to cover colouration and behavioural data from vast majority of the species included.

Taken together, our results are in accordance with previous studies, showing that aposematism is often associated evolutionarily with gregariousness. However, aposematism can also be lost while gregariousness persists, showing the traits are not obligately coupled. Extending the phylogeny to include close relatives of Diprionidae would help clarify the broader evolutionary picture of how gregariousness and aposematism have co-evolved in this lineage.

## Notes

### Competing Interest Statement

The authors have declared no competing interest.

### Funding

- Emil Aaltosen Säätiö
- Research Council of Finland

