## Supplementary Table 1 for "PHYLOGENETIC SUPPORT FOR THE EVOLUTION OF APOSEMATISM BEFORE GREGARIOUSNESS IN CONIFER SAWFLIES"

Table S1. Diprionid species in the phylogeny (n=58). Colouration data were available for 48 species and gregariousness data for 48 species. Data on both colouration and gregariousness were available for 45 species. A = aposematic, C = cryptic, G = gregarious, S = solitary and NA = no data available. In references, CRL = Catherine Linnen personal observation, MM = Marko Mutanen personal observation.

| **Species** | **Colouration** | **Behaviour** | **References for behaviour data** |
|---|---|---|---|
| Augomonoctenus smithi | NA | NA | NA |
| Diprion jingyanensis | NA | NA | NA |
| Diprion nanhuaensis | NA | G | MingCheng & Zheng 2010 |
| Diprion nipponicus | NA | NA | NA |
| Diprion pini | A | G | Hanski 1987, Lindstedt et al. 2011 |
| Diprion similis | A | S | CRL, MM |
| Gilpinia abieticola | C | S | Viitasaari & Varama 1987 |
| Gilpinia albiclavata | NA | NA | NA |
| Gilpinia frutetorum | C | S | Knerer & Atwood 1977, Kalin & Knerer 1977 |
| Gilpinia hercyniae | C | S | Knerer & Atwood 1977 |
| Gilpinia pallida | C | G | Hanski 1987, Viitasaari & Varama 1987 |
| Gilpinia polytoma | C | NA | NA |
| Gilpinia socia | A | G | Hanski 1987, Viitasaari & Varama 1987, Dobreva et al. 2022 |
| Gilpinia variegata | C | S | Hanski 1987, Viitasaari & Varama 1987 |
| Gilpinia virens | C | S | Hanski 1987, Viitasaari & Varama 1987 |
| Macrodiprion nemoralis | A | S | Hanski 1987, Viitasaari & Varama 1987, Zúbrik et al. 2013 |
| Microdiprion hakusanus | NA | NA | NA |
| Microdiprion pallipes | C | S | MM |
| Monoctenus juniperi | C | S | MM |
| Monoctenus obscuratus | A | S | MM |
| Neodiprion abbotii | C | G | Haack & Mattson 1993 |
| Neodiprion abietis | C | G | Haack & Mattson 1993, Davis et al. 2023 |
| Neodiprion autumnalis | A | G | McCullough & Wagner 1987, Ciesla & Smith 2011 |
| Neodiprion compar | C | S | Terbot et al. 2017, Davis et al. 2023 |
| Neodiprion dailingensis | NA | NA | NA |
| Neodiprion deleoni | C | NA | NA |
| Neodiprion demoides | C | G | CRL |
| Neodiprion dubiosus | A | G | Haack & Mattson 1993 |
| Neodiprion edulicolis | C | G | McGregor & Sandin 1969 |

| | | | |
|---|---|---|---|
| Neodiprion excitans | A | G | Bugwoodwiki, Thatcher 1971 |
| Neodiprion fabricii | C | G | CRL |
| Neodiprion gillettei | C | G | Ciesla & Smith 2011 |
| Neodiprion hetricki | A | G | CRL |
| Neodiprion knereri | A | G | USDA 2020, CRL |
| Neodiprion lecontei | A | G | Kalin & Knerer 1977, Terbot et al. 2017 |
| Neodiprion maurus | A | G | Haack & Mattson 1993, Terbot et al. 2017 |
| Neodiprion merkeli | C | G | CRL |
| Neodiprion mundus | C | G | Ciesla & Smith 2011 |
| Neodiprion nanulus | C | G | Haack & Mattson 1993 |
| Neodiprion nigroscutum | C | G | Becker & Benjamin 1967, CRL |
| Neodiprion omosus | C | G | Castañeda-Vildózola et al. 2021 |
| Neodiprion pinetum | A | G | Rauf & Benjamin 1980, Haack & Mattson 1993 |
| Neodiprion pinusrigidae | A | G | CRL |
| Neodiprion pratti | A | G | Ghent 1960, Haack & Mattson 1993, Bugwoodwiki |
| Neodiprion rugifrons | A | G | Kalin & Knerer 1977, Haack & Mattson 1993 |
| Neodiprion scutellatus | C | NA | NA |
| Neodiprion sertifer | C | G | Eisner 1974, Hanski 1987, Lindstedt et al. 2011 |
| Neodiprion swainei | A | G | McLeod 1970, Haack & Mattson 1993, Bugwoodwiki |
| Neodiprion taedae | A | G | Coyne 1970, Bugwoodwiki |
| Neodiprion tsugae | C | G | USDA 2021, Bugwoodwiki |
| Neodiprion ventralis | A | G | Ciesla & Smith 2011 |
| Neodiprion virginiana | A | G | CRL |
| Neodiprion warreni | A | G | CRL |
| Neodiprion xiangyunicus | NA | G | Hengxiao et al. 1993 |
| Nesodiprion flavipes | NA | NA | NA |
| Nesodiprion kojimai | NA | G | Plazi |
| Zadiprion rohweri | A | G | CRL |
| Zadiprion townsendi | A | G | Ciesla & Smith 2011, Bugwoodwiki |

Bugwoodwiki (visited 9/2026):
Neodiprion exitans: wiki.bugwood.org/Neodiprion_excitans
Neodiprion pratti banksianae: wiki.bugwood.org/Archive:Northeast/Neodiprion_pratti_banksianae Neodiprion swainei: wiki.bugwood.org/Neodiprion_swainei

Neodiprion tsugae: wiki.bugwood.org/Neodiprion_tsugae
Neodiprion taedae linearis: wiki.bugwood.org/Neodiprion_taedae_linearis
Zadiprion townsendi: wiki.bugwood.org/HPIPM:Zadiprion_townsendi

Castañeda-Vildózola A, Sánchez-Pale JR, Franco-Mora O & Hernández-Zetina DA. 2021. First record in Mexico of Ceromasia auricaudata Townsend (Diptera: Tachinidae) parasitizing Neodiprion omosus Smith (Hymenoptera: Diprionidae). *Florida Entomologist,* 104: 2.

Ciesla WM & Smith DR. 2011. Diprionid sawflies on lodgepole and ponderosa pines. *United States Department of Agriculture Forest Insect & Disease Leaflet*, 179: 1–11.

Coyne JF. 1970. Neodiprion taedae linearis: A sawfly pest of loblolly and shortleaf pines. U.S. Department of Agriculture Forest Service, Forest Pest Leaflet 34.

Davis JS, Glover AN, Everson KM et al. 2023. Identification, biology, and management of conifer sawflies (Hymenoptera: Diprioninae) in eastern North America. *J Integr Pest Manag*,14:13.

Dobreva M, Zlatanov N & Georgiev G. 2022. Two new sawfly species for the Bulgarian fauna (Hymenoptera: Symphyta). *Historia naturalis bulgarica,* 44: 45–49.

Eisner T, Johnessee JS & Carrel J. 1974. Defensive use by an insect of a plant resin. *Science*, 184: 996–999.

Ghent AW. 1960. A Study of the Group-Feeding Behaviour of Larvae of the Jack Pine Sawfly, Neodiprion pratti banksianae Roh. *Behavbiour*, 16: 110–148.

Haack RA & Mattson WJ. 1993. Life History Patterns of North American Tree-Feeding Sawflies. In: Sawfly Life History Adaptations to Woody Plants. Eds: Wagner MR & Raffa KF. Academic Press.

Hanski I. 1987. Pine sawfly population dynamics: patterns, processes, problems. *Oikos*, 50: 327–335.

Hengxiao G, Kidd NAC, Zhou JH & Xu XQ. 1993. Spatial distribution of the early larvae and cocoons of the pine sawfly Neodiprion xiangyunicus (Xiao & Huang) (Hym., Diprionidae) in China, and determination of an optimal population sampling programme. *J Appl Ent*, 115: 277–2833.

Kalin M & Knerer G. 1977. Group and mass effects in diprionid sawflies. *Nature*, 267: 427–429.

Knerer G & Atwood C. 1973. Diprionid Sawflies: Polymorphism and Speciation. *Science,* 179: 1090–1099.

Lindstedt C, Huttunen H, Kakko M & Mappes J. 2011. Disentangling the evolution of weak warning signals: high detection risk and low production costs of chemical defences in gregarious pine sawfly larvae. *Ecol Evol*, 25: 1029–1046.

McCullough DG & Wagner MR. 1987. Influence of watering and trenching ponderosa pine on a pine sawfly. *Oecologia*, 71: 382–387.

McGregor MD & Sandin O. 1969. Pinyon Sawfly, *Neodiprion edulicolus* Ross. U.S. Department of Agriculture Forest Service, Forest Pest Leaflet 117.

McLeod JM. 1970. The Epidemiology of the Swaine Jack-pine Sawfly, Neodiprion Swainei Midd. The Forestry Chronicle April 1970. Canadian Forestry Service.

MingCheng J & Zheng P. 2010. Preliminary study on biological characteristics and control of Diprion nanhuaensis. *Journal of Jiangsu Forestry Science & Technology,* 37: 28–30.

Plazi: treatment.plazi.org/id/643687C2D85AFFBEFF7BFCAAF263F953/12 (visited 8/2025).

Rauf A & Benjamin DM. 1980. The Biology of the White Pine Sawfly, Neodiprion pinetum (Hymenoptera: Diprionidae) in Wisconsin. *The Great Lakes Entomologist*, 13: 4.

Terbot JW, Gaynor RL & Linnen CR. 2017. Gregariousness does not vary with geography, developmental stage, or group relatedness in feeding redheaded pine sawfly larvae. *Ecology and Evolution*, 7: 3689–3702.

Thatcher RC. 1971. *Pine Sawfly* Neodiprion excitans *Roh*. U.S. Department of Agriculture Forest Service, Forest Pest Leaflet 105.

USDA 2020: Florida Forest Service Forest Health Highlights. 2020. U.S. Department of Agriculture, Forest Service. 2020.

USDA 2021. Forest Insect and Disease Leaflet 31. Forest Service U.S. Department of Agriculture. 2021.

Viitasaari M & Varama M. 1987. Sahapistiäiset 4. University of Helsinki, Department of Agricultural and Forest Zoology, Reports 10, Helsinki.

Zúbrik M, Kunca A & Csóka G. 2013. Insects and diseases damaging trees and shrubs of Europe. NAP Editions.
